# Diversity loss selects for altered plant phenotypic responses to local arbuscular mycorrhizal communities

**DOI:** 10.1101/216986

**Authors:** Terhi Hahl, Sofia J. van Moorsel, Marc W. Schmid, Debra Zuppinger-Dingley, Bernhard Schmid, Cameron Wagg

**Affiliations:** URPP Global Change and Biodiversity and Department of Evolutionary Biology and Environmental Studies, University of Zürich, Winterthurerstrasse 190, CH-8057, Switzerland; MWSchmid GmbH, Möhrlistrasse 25, CH-8006 Zürich

**Author notes:** Equal contribution (Terhi Hahl, Sofia J. van Moorsel). Author for correspondence: Cameron Wagg.

**Keywords:** biodiversity experiment, functional traits, growth–defence trade-off, mutualism–parasitism continuum, plant–AMF interactions, plant–soil feedbacks

## Abstract

1. Biodiversity loss not only impairs ecosystem functioning but can also alter the selection for traits in plant communities. At high diversity selection favours traits that allow for greater niche partitioning, whereas at low diversity selection may favour greater defence against pathogens. However, it is unknown whether changes in plant diversity also select for altered interactions with soil organisms.
2. We assessed whether the responses in plant growth and functional traits to their local arbuscular mycorrhizal fungal (AMF) communities have been altered by the diversity of the plant communities from which both plants and AMF communities were obtained. We grew plants with AMF communities that originated from either plant monocultures or mixtures in a fully factorial design that included both negative and positive controls, by inoculating no AMF or a foreign AMF respectively.
3. We found that AMF from plant mixtures were more beneficial than monoculture AMF for two out of five plant species. Plants from mixtures generally grew better than those from monocultures, but suffered greater damage by leaf pathogens. Although plant growth and phenotypic responses were dependent on the AMF communities with which they associated, we found little evidence for plant growth responses specific to their local AMF communities and results differed between species and traits.
4. Our results show that plants from mixtures were selected for increased growth at the expense of reduced defence and vice versa for plants from monocultures, providing evidence for plant diversity-dependent selection on competitive growth vs. defence. Furthermore, our study suggests that effects of a common history between plants and AMF do not follow a general pattern leading to increased or decreased mutualism.
5. *Synthesis:* Here we provide evidence that biodiversity loss can alter evolutionary trajectories of plant phenotypes and responses to their local AMF communities. However, the selection for altered plant–AMF interactions differ between plant species. To understand how plant communities respond and evolve under a changing environment requires further knowledge about life strategies of plant species and their above–belowground interactions.

## 1 Introduction

How reciprocal interactions between plants and soil biota drive plant community structure and ecosystem functioning has been a focal point in ecology (Bever, 1994; Klironomos, 2002; van der Heijden et al., 2006; van der Putten et al., 2013). These plant–soil feedbacks (PSF) describe the relationships between plants and their associated soil pathogens and mutualists, which negatively or positively affect plant performance (Bever et al., 2010). Negative PSF are common and thought to be key to maintaining diversity in plant communities and to successional changes in plant community composition through time (Kardol, Martijn Bezemer, & van der Putten, 2006; Klironomos, 2002; Andrew Kulmatiski, Beard, Stevens, & Cobbold, 2008; Petermann, Fergus, Turnbull, & Schmid, 2008; van der Putten et al., 2013). Positive PSFs are known to improve plant performance by either enhancing pathogen defence (reducing negative PSF) or stimulating plant growth (van der Putten et al., 2013). Plants simultaneously interact with both pathogens and beneficial soil organisms that together determine plant performance. However, negative PSF and positive PSF have rarely been disentangled and only recently studies considered that both negative and positive PSF might change over ecological time-scales (van der Heijden, Bardgett, & van Straalen, 2008; terHorst et al. 2014). Here we assess the consequences of plant diversity loss on interactions between plants and associated communities of soil mutualists. We specifically tested whether changes in plant diversity result in intraspecific differences in plant phenotypic responses to their local arbuscular mycorrhizal fungi (AMF) communities over a short evolutionary time scale (11 years).

Arbuscular mycorrhizal fungi (AMF) are soil-borne fungi, which form symbiotic relationships with most land plants (Harley & Harley, 1987; Smith & Read, 1997; Wang & Qiu, 2006). These fungi are typically known for improving plant growth by providing enhanced soil nutrient uptake in exchange for plant photosynthates (Smith & Read 2008). Furthermore, AMF diversity has been shown to be an important driver of aboveground plant diversity and ecosystem productivity (Francis & Read, 1994; van der Heijden et al., 1998; Wagg, Jansa, Stadler, Schmid, & van der Heijden., 2011a). Conversely, the composition of AMF communities can be strongly affected by the composition and diversity of plant communities, such that the diversity of plant communities is interlinked with the diversity of AMF communities (Burrows & Pfleger, 2002; Klironomos, McCune, Hart, & Neville, 2000; König et al., 2010; Milcu et al., 2013; Scherber et al., 2010). It is therefore conceivable that a more diverse plant community can sustain a more beneficial AMF community due to a positive effect of plant diversity on supporting a greater AMF diversity. On the other hand, plants growing in monocultures may also select for a more beneficial AMF community, as the greater abundance of a specific plant species could select for AMF that are particularly beneficial to that particular species (Arguello et al., 2016; Kiers et al., 2011; Werner & Kiers, 2015). Although there is evidence indicating that plants growing in monoculture for a prolonged period of time had evolved altered phenotypic responses to competition and positive PSF (Zuppinger-Dingley et al., 2014; Zuppinger-Dingley, Flynn, De Deyn, Petermann, & Schmid, 2016), we currently lack evidence as to whether changes in plant diversity alter plant responses to their local AMF communities over time. We therefore test the hypothesis that AMF communities originating from plant communities differing in diversity (plant monocultures vs. plant mixtures) would differ in their influence on plant growth and phenotypic traits (H1).

Plants can adapt to short-term selection pressure imposed by plant community diversity (van Moorsel, Schmid, Hahl, Zuppinger-Dingley, & Schmid, 2017; Zuppinger-Dingley et al., 2014). This rapid evolution in plant communities can occur via a sorting-out from standing genetic variation (Fakheran et al., 2010). In monocultures, plants may accumulate more specialized pathogens than in diverse plant communities, which potentially reduces plant productivity in monocultures over time (Kulmatiski, Beard, & Heavilin, 2012; Marquard et al., 2013; Schnitzer et al., 2011; van der Putten et al., 2013). At higher plant diversity levels, however, specialized pathogens may become diluted (Eisenhauer, 2012; Latz et al., 2012), thus conferring lower pathogen-based selection pressure than in monocultures. Consequently, in plant mixtures, plants would likely trade-off reduced defence for increased competitive growth, whereas in monocultures plants would trade-off reduced growth for increased defence (Herms & Mattson, 1992). If plants are differentially selected for pathogen defence or competitive growth traits depending upon their neighbourhood diversity from which they originate (e.g. Zuppinger-Dingley et al. 2014, Zuppinger-Dingley et al. 2016), then we hypothesize that plants with different selection backgrounds exhibit different phenotypic traits related to competition and pathogens (H2).

Furthermore, in plant monocultures under a high pathogen load (Schnitzer et al., 2011), it may be particularly important for plants to associate with AMF as these fungi are known to protect plants against pathogens (Newsham, Fitter, & Watkinson, 1995; Pozo & Azcón-Aguilar, 2007). The increasing positive PSF on monocultures through time (e.g. Zuppinger-Dingley et al. 2016) could thus be potentially reflecting co-selection of plant–AMF relationships. Considering the potential for plant diversity-driven trade-offs between competitive growth and plant defence, it is likely that over time plants and their associated soil biota, such as AMF, may have altered PSF depending on the diversity of the plant community from which they have originated. We therefore finally hypothesize that differences in plant diversity have differentially altered plant phenotypic responses to their local AMF communities resulting in specific ‘home’ vs. ‘away’ plant–AMF community interactions (H3). Home vs. away effects have been used in reciprocal transplant experiments (Joshi et al., 2001) and reciprocal inoculation experiments (Klironomos, 2002; Wagg, Husband, Massicotte, & Peterson, 2011b) to describe a matching of locally co-occurring populations. In the present study, we compared home-combinations (same history for both AMF and plants) with away-combinations (different history for the two partners).

To address our three hypotheses, we used five plant species that occurred in plant mixtures and as monocultures within the Jena Biodiversity Experiment (Roscher et al., 2004) and the AMF spore communities collected from their rhizospheres. Plants and AMF had thus been co-occurring in natural field conditions for eleven years (van Moorsel et al., 2018). To specifically address the home and away plant–AMF interactions, we isolated AMF spore communities from the rhizosphere soil and propagated them in trap cultures. Our design was a reciprocal inoculation experiment where plants with a history of occurring in monocultures (monoculture-type plants) or diverse plant mixtures (mixture-type plants) were inoculated with AMF communities isolated from the same monocultures (monoculture AMF) or mixtures (mixture AMF). In addition, we included a negative AMF control treatment (no AMF present) as well as a positive control where plants were inoculated with an external AMF species *(Rhizoglomus irregulare* (Blaszk., Wubet, Renker & Buscot) Sieverd., G.A. Silva & Oehl), which did not share any history with the plants.

## 2 Materials and methods

### 2.1 PLANT SELECTION HISTORIES

We used five common perennial European grassland species from three functional groups (Roscher et al., 2004): three small herbs *(Plantago lanceolata* L., *Prunella vulgaris* L. and *Veronica chamaedrys* L.), one tall herb *(Galium mollugo* L.) and one legume *(Lathyrus pratensis* L.). The plants had a selection history from the Jena Experiment (Roscher et al., 2004), where they were sown in either monoculture or mixture in 2002 (selection history “monoculture-type plants” and “mixture-type plants”, respectively). The species compositions in the experimental plots in Jena were maintained by weeding three times per year in spring, summer and autumn and by mowing twice per year at peak biomass times in spring and summer. Each of the studied plant species had undergone eleven years of selection from 2002 until 2014 in either plant monocultures (monoculture-type plants) or mixtures (mixture-type plants, Fig. 1). The five plant species are known to associate with arbuscular mycorrhizal fungi (Harley & Harley, 1987).

**Figure 1.**
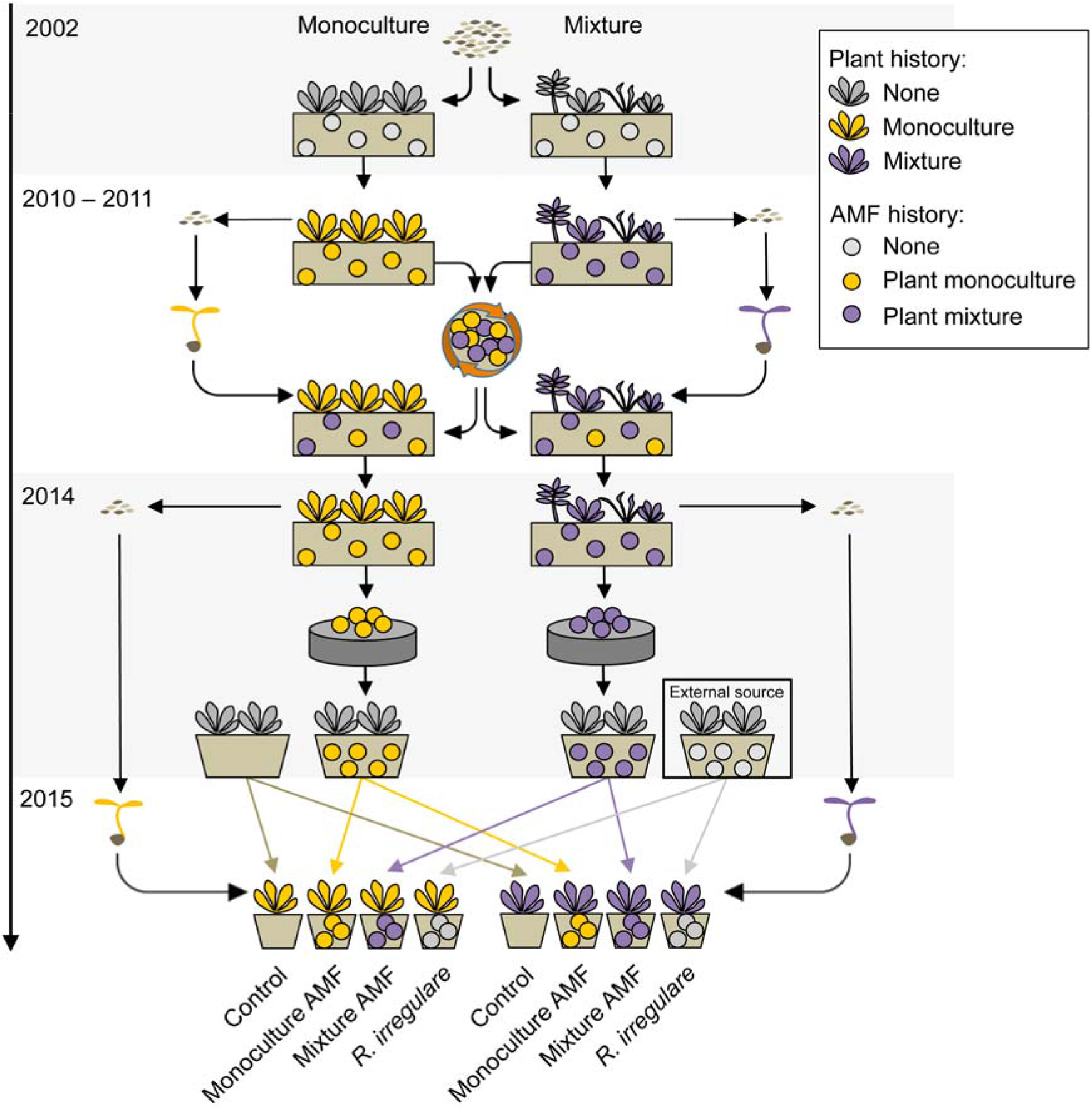
Experimental design. Plant monocultures and mixtures in the Jena Experiment were sown in 2002 and maintained until 2010. In 2010, the plants of 48 plots underwent a first controlled seed production event and the soil of the plots was pooled, mixed and placed back to the excavated locations. In spring 2011, the seedlings produced were transplanted back to the mixed soil in the same plots from which their parents were excavated. The plant communities could then again associate with their own microbial communities potentially co-assembling and co-evolving until 2014. In spring 2014, the plants underwent a second controlled seed production event, and the AMF spores from their rhizosphere soil were isolated. The isolated AMF communities accumulated in trap-cultures for ten months with trap plants lacking a common selection history with the AMF spores. Control trap-cultures without AMF spores were established as negative control. Four treatments were created: pots with sterile soil and 1) 9 % inoculum without AMF, 2) inoculum of AMF isolated from plants grown in monoculture, 3) inoculum of AMF isolated from plants grown in mixture and 4) inoculum containing *Rhizoglomus irregulare.* Finally, the plants with a selection history in either monoculture (monoculture-type plants) or mixture (mixture-type plants) were planted individually into the prepared pots.

### 2.2 FIRST CONTROLLED SEED PRODUCTION

In spring 2010, entire plant communities of 48 plots (12 monocultures, 12 two-species mixtures, 12 four-species mixtures and 12 eight-species mixtures) of the biodiversity experiment in Jena, Germany (the Jena Experiment) were collected as cuttings. Additionally, the top 30 cm soil of the 48 plots was pooled together, mixed and placed back into the excavated locations at the Jena Experiment. The cuttings were then transplanted to an experimental garden in Zurich, Switzerland, in identical plant composition and seeds were collected over summer 2010. The plots were filled with 30 cm of soil (1:1 mixture of garden compost and agricultural field soil, pH 7.4, Gartenhumus, RICOTER Erdaufbereitung AG, Aarberg, Switzerland), and fenced with netting to minimize cross-pollination with plants outside the plots (for details see Zuppinger-Dingley et al. 2014). In spring 2011, seedlings were propagated from the collected seeds in a glasshouse in Zurich and then transplanted back into the mixed soil in the same plots of the Jena Experiment from where their parents had originally been excavated. In the re-established plots, these newly established plant communities were maintained for three years until 2014 to allow them to become associated with their own microbial communities (van Moorsel et al., 2018).

### >2.3 SEED PRODUCTION AND SOIL COLLECTION

In March 2014, entire plant communities from the re-established plots at the Jena Experiment were collected and planted in their respective communities in 1 x 1 m plots in the experimental garden in Zurich. The plots were filled with 30 cm of soil (1: 1 mixture of garden compost and agricultural field soil, pH 7.4, Gartenhumus, RICOTER Erdaufbereitung AG, Aarberg, Switzerland), and fenced with netting to minimize cross-pollination. During the relocation of the plant communities, we collected rhizosphere soil samples attached to the roots of the plants (Fig. 1). By then, the soil communities had undergone three years of establishment and eight plus three years of potential co-selection with each of the five plant species in monocultures or mixtures. Seeds were collected from five monocultures and five mixtures (one four-species mixture and four eight-species mixtures). The seeds of the five plant species were stored at +4 °C for two months. Four weeks before the start of the experiment, the seeds were surface-sterilized in 7–14 % bleach for 10–45 min and subsequently germinated on 1% water-agar.

### 2.4 INOCULUM PREPARATION

We created trap cultures using AMF spore isolation to minimize the potential that other unintended fungi were also propagated. To isolate AMF spore communities from the sampled rhizosphere soils, we passed deionized water and 25 g of soil sample through a series of sieves from 500 μm to 32 μm using a sugar gradient-centrifugation method (Sieverding, 1991). The AMF spores were manually collected with a pipet under a microscope at 200-fold magnification. To bulk up the isolated AMF communities for use as inoculum, we established trap cultures that consisted of 2 L of 4:1 sand-soil mixture, autoclaved at 120 °C for 99 min, and a monoculture of trap plants of one the five tested plant species (Fig. 1). All trap cultures received approximately 400 AMF spores in 30 ml of deionized water, except for the negative control trap cultures, which received 30 ml of deionized water without AMF spores. For the trap-culture plants we used new seeds from a commercial seed supplier that provided the original seed material for the Jena Experiment (Rieger-Hofmann GmbH, Blaufelden-Raboldshausen, Germany). The seeds were surface-sterilized in 7–14 % bleach for 10–45 min and pre-germinated on 1 % water agar. Each AMF trap culture was replicated twice. After ten months of growth in the glasshouse, we collected root samples from each trap culture that were then fixed in 50 % ethanol, cleared in 10 % KOH, and stained in 5 % ink-vinegar (Vierheilig, Coughlan, Wyss, & Piché, 1998). Colonization by AMF was microscopically assessed for successful mycorrhizal establishment by quantifying AMF root colonization following the transect-intersect method (McGonigle, Miller, Evans, Fairchild, & Swan, 1990). We further quantified the concentration of AMF spores in trap cultures with fungal colonization. We isolated AMF spores from a 10-g soil sample with the same sieving and centrifugation methods used when setting up the AMF trap-culture pots. Trap-plant cultures that exhibited fungal root colonization and spore production were dried and the plants were harvested at ground level. The roots were harvested, cut into 3–5 cm fragments and the belowground content of the trap cultures was used as soil inoculum in the PSF experiment described below.

For the positive control AMF treatment, we used a trap culture substrate containing *Rhizoglomus irregulare* (Blaszk., Wubet, Renker & Buscot) Sieverd., G.A. Silva & Oehl as the inoculum. *Rhizoglomus irregulare* (previous names *Glomus intraradices* and *R. irregularis*; (Sieverding, da Silva, Berndt, & Oehl, 2015) is an AMF taxon common in natural grasslands. The culture was developed for nine months in a substrate of 15 % soil, 65 % sand and 20 % oil binder with *P. lanceolata* plants, which had no shared selection history with plants or soils from the Jena Experiment. The *R. irregulare* material was obtained from M.G.A. van der Heijden’s Ecological Farming Group (Agroscope Reckenholz-Tänikon, Zurich, Switzerland).

### 2.5 EXPERIMENTAL DESIGN

To establish the AMF treatments, we filled 1-L pots with gamma-radiated (27–54 kGy) 1:1 (weight/weight) sand-soil mixture and added 9 % (volume/volume) of inoculum without AMF (negative control), inoculum of AMF isolated from plants grown in monoculture (monoculture AMF) or mixture (mixture AMF) or inoculum containing *R. irregular* (a positive control). One monoculture- or mixture-type plant of a single plant species was planted into each pot (Fig. 1, lower panel). To standardize the non-AMF microbial community within each pot, we created a microbial wash by filtering 1.2 L of a mixture of unsterilized field soil and the AMF trap culture substrates through a series of sieves and finally through filter paper (MN615, Macherey-Nagel GmbH & Co. KG) with 5 L of deionized water. Each pot received 10 ml of the microbial-wash filtrate. The experiment included four AMF treatments in total, two plant histories (monoculture- and mixture-type plants) and five plant species in a full factorial design (Table S1). Control and *R. irregulare* AMF treatments were replicated five times and the two other AMF treatments were replicated ten times (five times per trap-culture replicate, Table S1). For the mixture-type plants of *G. mollugo*, we did not have sufficient seedlings for the full design and the AMF treatments were thus only replicated 9 and 8 times, respectively (Table S1). The 297 pots were randomly arranged within five experimental blocks in a glasshouse compartment with each particular treatment combination and trap-culture replicate occurring only once in each block.

### 2.6 DATA COLLECTION

We cut the plants to 4 cm aboveground three months after planting seedlings into the pots of the AMF treatments (referred to as first harvest). After five months of plant growth, maximum height and average leaf absorbance (SPAD-502Plus Chlorophyll Meter, KONICA MINOLTA, INC., Osaka, Japan) of three representative leaves of each plant were measured and the aboveground biomass was harvested at ground-level (referred to as second harvest). The biomass of each plant was dried at 70 °C for 48 h and then weighed. We assessed leaf mass per area (LMA) and leaf dry matter content (LDMC) at the second harvest by measuring the area of fresh leaves (LI-3100C Area Meter, LI-COR, Lincoln, USA) immediately after harvest and assessing the weight of the leaves before (fresh weight) and after drying (dry weight). Finally, we estimated the degree of damage on plant aboveground tissues due to powdery mildew (family Erysiphaceae) and two-spotted spider mites *(Tetranychus urticae* Koch). To determine AMF colonization, roots were washed free of adhering rhizosphere soil and cut into small 1–5 cm fragments and a random subsample of roots were then stored in 50 % ethanol for microscopic quantification of AMF using the same clearing, staining and scoring methods as described above. All measured traits are listed in Table S2.

### 2.7 DATA ANALYSES

Due to a contamination of control soil with AMF, one pot with *L. pratensis* was excluded from all analyses. The biomass data, morphological trait measurements, leaf damage estimates and AMF colonization were analysed using linear models. Because the first measure assessed growth and the second regrowth, the harvests were analyzed separately. Plant survival and AMF presence/absence were analysed using analysis of deviance. The results were summarized in analysis of variance (ANOVA) and deviance (ANDEV) tables (McCullagh & Nelder, 1998; Schmid, Baruffol, Wang, & Niklaus, 2017). The explanatory terms of the models were block, plant history (monoculture-type vs. mixture-type), AMF treatments (four AMF treatments or sequence of the following three orthogonal contrasts: control vs. AMF treatments, *R. irregulare* vs. monoculture or mixture AMF and monoculture vs. mixture AMF), species and interactions of these. Statistical analyses were conducted using the software product R, version 3.0.2 (R Core Team, 2013).

## 3 Results

### 3.1 ROOT COLONIZATION

The root colonization by AMF was highest in all five plant species with the inculcation of *R. irregulare* (the positive control). For two species, *G. mollugo* and *L. pratensis*, AMF colonization was higher in the treatment with AMF originating from the plant monocultures, whereas for two other species, *P. lanceolata* and *V. chamaedrys*, root colonization was greater with AMF originating from plant mixtures. In *P. vulgaris*, AMF colonization was equally low for both AMF community histories. An increase in AMF colonization was positively correlated with an increase in individual plant biomass for all three AMF treatments (Fig. S1).

### 3.2 EFFECTS OF AMF COMMUNITIES ON PLANT PHENOTYPIC RESPONSES (H1)

The four AMF treatments changed the phenotypes of the plants, but the response varied between the five species. We found significant effects of the AMF treatments on five traits: plant aboveground biomass at the first harvest, plant aboveground biomass at the second harvest, plant height, LDMC and AMF colonization (Fig. 2, Table 1). The effects of the AMF treatments on plant biomass were relatively consistent between the two harvests (Fig. 2). Biomass was generally highest in the treatment with the positive AMF control (inoculation of *R. irregulare*), with the exception of *V. chamaedrys*, for which the negative control (no AMF inoculated) resulted in the highest biomass. Soil containing AMF from mixture plots was the overall second most productive AMF treatment in terms of plant biomass, but again with the exception of *V. chamaedrys* and also *P. lanceolata* which exhibited higher biomass when inoculated with AMF from monoculture plots. The effects of the AMF treatments on plant height also varied among plant species. For *G. mollugo*, soil containing *R. irregulare* or soil containing mixture AMF resulted in the tallest plants. For three species, *L. pratensis, P. lanceolata* and *P. vulgaris*, the AMF treatment did not have a strong effect on plant height, except that the control treatment lacking AMF resulted in the shortest plants. In line with the biomass results, plant height was increased in the control treatment for the small herb *V. chamaedrys*. The LDMC was strongly increased in the control treatment for *L. pratensis* and slightly increased for *G. mollugo.* For the three small herbs, *P. lanceolata, P. vulgaris* and *V. chamaedrys*, the LDMC was increased for the two AMF treatments containing AMF from either mixture or monoculture plots as opposed to treatments without AMF or with the external *R. irregulare.*

**Table 1.** ANOVA results for eight traits and ANDEV results for plant survival in response to the design variables.

|  | Plant survival |  |  | Biomass<br>first harvest |  |  | Biomass<br>second harvest |  |  | Plant height |  |  | LMA |  |  | LDMC |  |  | Leaf damage |  |  | Leaf absorbance |  |  | AMF colonization |  |  |
| --- | --- | --- | --- | --- | --- | --- | --- | --- | --- | --- | --- | --- | --- | --- | --- | --- | --- | --- | --- | --- | --- | --- | --- | --- | --- | --- | --- |
| Source of variation | *Df* | % Dev | *P* | *Df* | % SS | *P* | *Df* | % SS | *P* | *Df* | % SS | *P* | *Df* | % SS | *P* | *Df* | % SS | *P* | *Df* | % SS | *P* | *Df* | % SS | *P* | *Df* | % SS | *P* |
| Block | 4 | 0.81 | 0.448 | 4 | 6.25 | <0.001 | 4 | 6.63 | <0.001 | 4 | 1.31 | 0.018 | 4 | 4.85 | <0.001 | 4 | 5.58 | 0.002 | 4 | 5.68 | <0.001 | 4 | 3.40 | <0.001 | 4 | 1.70 | 0.130 |
| Plant history (PH) | 1 | 0.03 | 0.689 | 1 | 0.03 | 0.656 | 1 | 0.47 | 0.050 | 1 | 2.26 | <0.001 | 1 | 0.88 | 0.014 | 1 | 0.63 | 0.164 | 1 | 1.43 | 0.016 | 1 | 0.39 | 0.032 | 1 | 0.10 | 0.510 |
| AMF treatment (ST) | 3 | 1.44 | 0.088 | 3 | 3.52 | <0.001 | 3 | 0.67 | 0.139 | 3 | 0.44 | 0.258 | 3 | 0.95 | 0.090 | 3 | 0.25 | 0.852 | 3 | 0.57 | 0.503 | 3 | 1.58 | 0.000 | 3 | 38.21 | <0.001 |
| Species (Sp) | 4 | 31.48 | <0.001 | 4 | 45.11 | <0.001 | 4 | 62.74 | <0.001 | 4 | 69.90 | <0.001 | 4 | 59.82 | <0.001 | 4 | 20.53 | <0.001 | 4 | 35.70 | <0.001 | 4 | 75.64 | <0.001 | 4 | 7.21 | <0.001 |
| PH x ST | 3 | 0.34 | 0.668 | 3 | 0.06 | 0.935 | 3 | 0.20 | 0.646 | 3 | 0.38 | 0.315 | 3 | 0.47 | 0.357 | 3 | 0.72 | 0.524 | 3 | 0.04 | 0.982 | 3 | 0.04 | 0.916 | 3 | 0.11 | 0.923 |
| PH x Sp | 4 | 5.06 | <0.001 | 4 | 3.58 | <0.001 | 4 | 2.55 | <0.001 | 4 | 1.67 | 0.005 | 4 | 1.60 | 0.028 | 4 | 3.70 | 0.024 | 4 | 6.70 | <0.001 | 4 | 0.31 | 0.446 | 4 | 0.56 | 0.665 |
| ST x Sp | 12 | 3.85 | 0.131 | 12 | 4.44 | 0.005 | 12 | 2.94 | 0.024 | 12 | 3.37 | 0.003 | 12 | 2.18 | 0.245 | 12 | 7.09 | 0.045 | 12 | 3.07 | 0.396 | 12 | 1.46 | 0.147 | 12 | 9.02 | <0.001 |
| PH x ST x Sp | 12 | 2.35 | 0.546 | 12 | 3.00 | 0.081 | 11 | 1.73 | 0.224 | 11 | 1.03 | 0.568 | 11 | 3.30 | 0.024 | 11 | 3.61 | 0.429 | 11 | 2.59 | 0.470 | 11 | 1.78 | 0.038 | 11 | 2.00 | 0.671 |
| Residuals | 252 | 54.65 |  | 224 | 34.01 |  | 183 | 22.06 |  | 183 | 19.63 |  | 180 | 25.94 |  | 180 | 57.88 |  | 183 | 44.21 |  | 183 | 15.38 |  | 174 | 41.08 |  |

**Figure 2.**
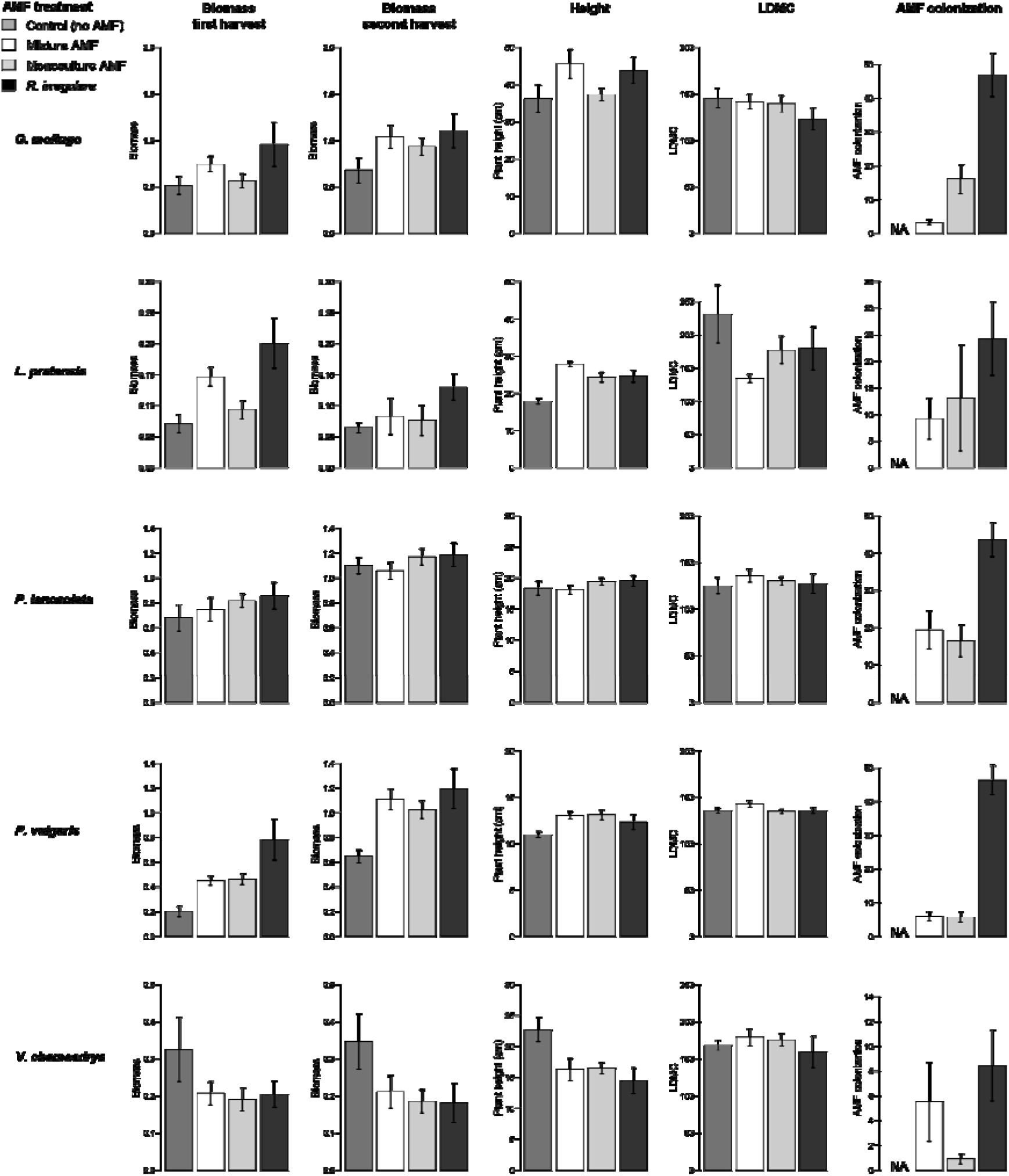
Functional traits in responses to the AMF treatments are shown separately for each of the five plant species assessed. Bars are means with standard errors calculated from raw data. See Table 1 for significance of effects.

### 3.3 INFLUENCE OF PLANT DIVERSITY HISTORY ON PLANT PHENOTYPES (H2)

Mixture-type plants grew taller than monoculture-type plants in three out of five species. Monoculture-type plants of the other two species, *L. pratensis* and of *P. lanceolata,* grew taller than their mixture-type counterparts. The leaf traits LDMC and LMA were generally increased in mixture-type plants, except for *V. chamaedrys.* For plant height, LMA, leaf damage and leaf absorbance the plant history effect was also visible across species (Table 1, e.g. the main effect of plant history).

As anticipated, mixture-type plants overall had more damaged leaves than monoculture-type plants, in particular for *L. pratensis* and *P. lanceolata* (Fig. 3, Table 1). The monoculture-type or mixture-type plants resulted in differential phenotypes between individuals of the same species. However, the strength of the response to selection history varied greatly between the five studied species (Fig. 3, Table 1). We found significant overall effects of plant history on seven traits: plant survival, plant aboveground biomass at the first harvest, plant aboveground biomass at the second harvest, plant height, LMA, LDMC and leaf damage. Mixture-type plants had a greater survival in the case of the three small herbs, whereas for the legume *L. pratensis* and the tall herb *G. mollugo*, monoculture-type plants showed a higher survival (Fig. 3). Plant biomass was generally higher for mixture-type plants (Fig. 3), but *L. pratensis* and *G. mollugo* showed the opposite pattern with increased biomass for monoculture-type plants (for *L. pratensis* only at the first harvest). The difference in biomass production between monoculture-and mixture-type plants was smaller at the second harvest but still varied significantly among species (Fig. 3).

**Figure 3.**
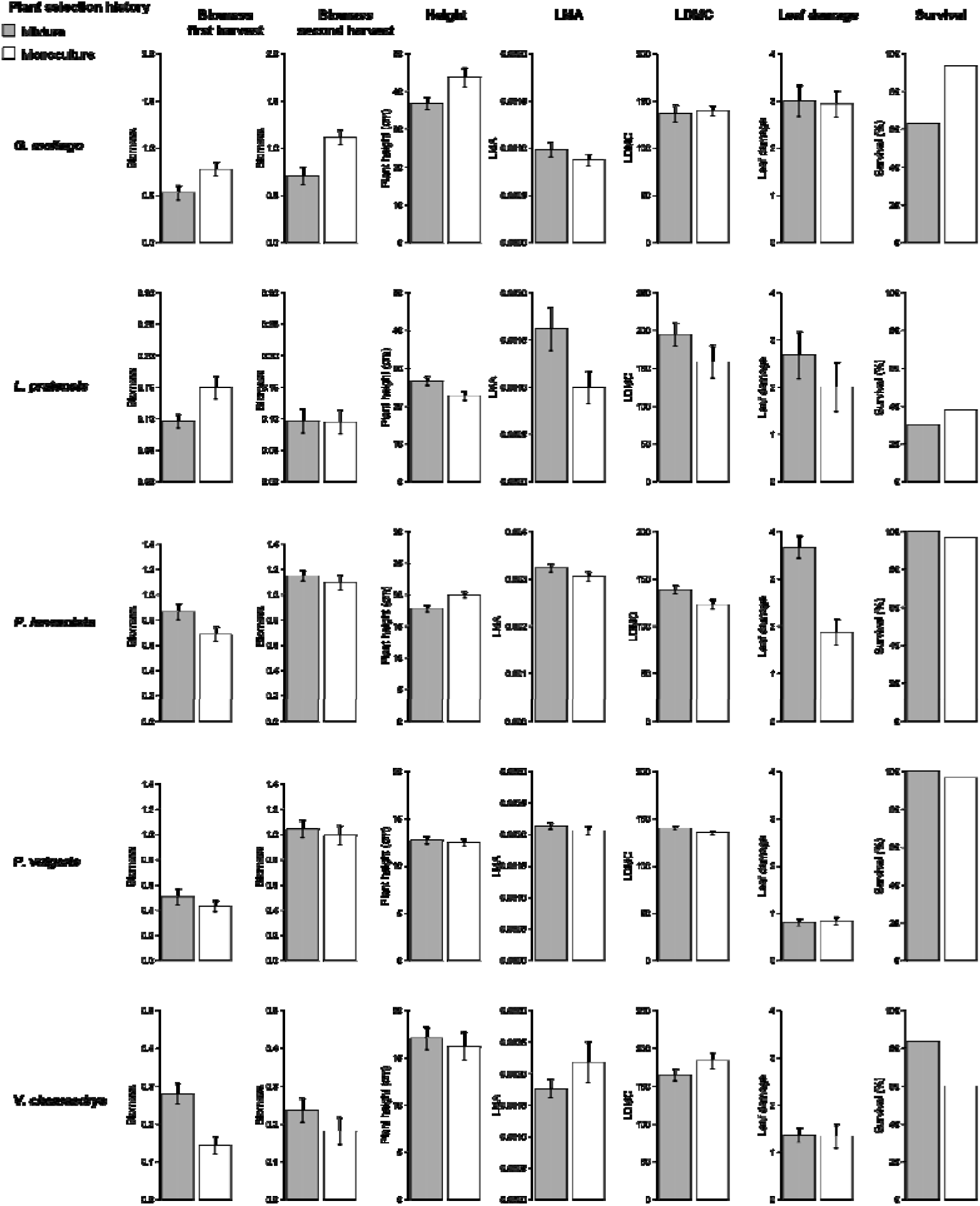
Functional traits in response to plant selection history for five species. Grey bars refer to plants with a history of growing in species mixtures at the Jena Experiment, white bars refers to plants with a history of growing in monoculture at the same field site. Shown are means and standard errors calculated from raw data. See Table 1 for significance of effects.

### 3.4 EFFECTS OF MATCHING HOME AND AWAY PLANT AND AMF HISTORIES (H3)

We did find overall home vs. away effects for LMA and leaf absorbance, but the effects varied among the five plant species studied and responses differed in direction (Fig. 4). The home mixture combination (mixture-type plants and mixture AMF) resulted in increased LMA in four out of five species, the exception being *V. chamaedrys* (Fig. 4a). The home monoculture combination (monoculture-type plants and monoculture-AMF) only increased LMA in *P. vulgaris* (Fig. 4a). Both away-combinations (monoculture-plant type with mixture AMF and vice versa) increased LMA strongly in *V. chamaedrys* (Fig. 4a). The away-combination between mixture-type plants and monoculture AMF also increased LMA in *L. pratensis* (Fig. 4a). Both home-combinations increased leaf absorbance in the small herbs *P. vulgaris* and *V. chamaedrys* (Fig. 4b). In contrast, both home-combinations reduced leaf absorbance for the tall herb *G. mollugo* and the legume *L. pratensis.*

**Figure 4.**
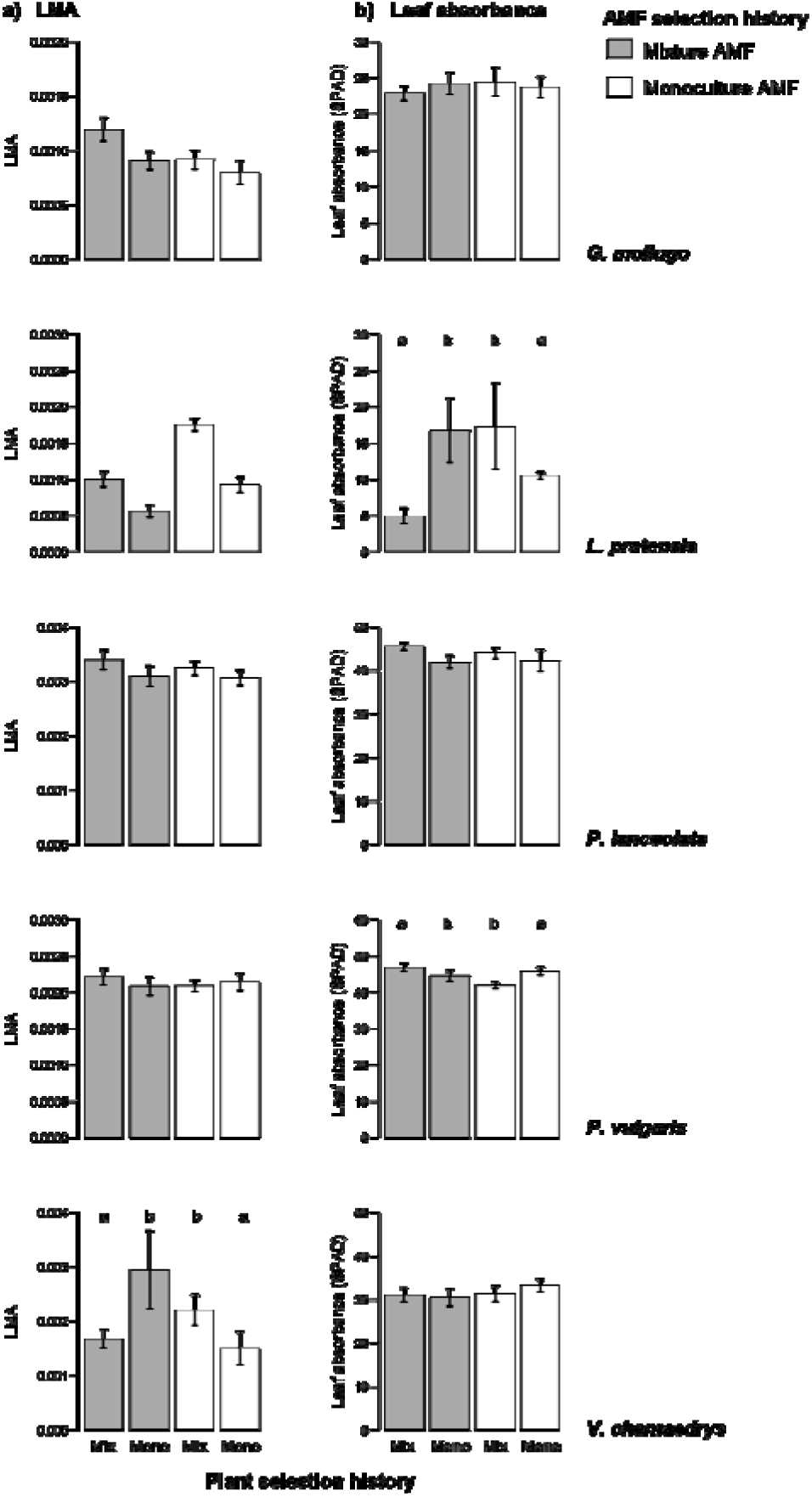
(a) LMA and (b) leaf absorbance in response to home and away combinations of plant selection history and AMF selection history for five species. Grey bars refer to AMF from mixed plant communities; white bars refer to AMF from monocultures. The first and last bar of each plot corresponds to home combinations; the two middle bars correspond to away combinations. Shown are means and standard errors calculated from raw data. Significant home effects are indicated by letters “a” and “b” on top of bars: *P* < 0.01 for leaf absorbance in *L. pratense, P* < 0.05 for leaf absorbance in *P. vulgaris* and *P* < 0.05 for LMA in *V. chamaedrys.*

## 4 Discussion

### 4.1. EFFECTS OF AMF COMMUNITIES ON PLANT PHENOTYPIC RESPONSES (H1)

Here we assessed whether plant diversity loss not only alters plant traits related to competitition, but also the phenotypic responses of plants to their AMF communities. Specifically, we aimed to separate whether plant diversity loss alters the selection on plants for altered responses to local AMF communities (H1), traits related to competitive growth (H2) or both simultaneously (H3). For instance, although plants may be selected for altered competitive growth characteristics in a more species diverse community (Zuppinger-Dingley et al. 2014), it is also known that interactions with soil organisms alters how plants allocate resources to traits, which affects plant growth strategies (Dudenhöffer, Ebeling, Klein, & Wagg, 2018; Streitwolf-Engel et al., 1997). Conversely, plant traits have been thought to influence interactions between plants and AMF (Baxendale, Orwin, Poly, Pommier, & Bardgett, 2014). A recent study focusing on plant traits in relation to PSF effects found evidence that a lower specific root length and a greater percentage colonization of the root length by AMF resulted in more positive PSFs (Cortois et al. 2016). Furthermore, mycorrhizal associations have been known to alter leaf chemical properties and alter herbivore life-history traits via changes in leaf chemistry (Goverde, van der Heijden, Wiemken, Sanders, & Erhardt, 2000; Smith & Read, 1997).

We assessed whether plant diversity can act as a selective environment on how AMF communities influence plant growth and functional traits. In support of our first hypothesis (H1) we found significant effects of the AMF treatments on plant phenotypes. Mixture AMF were more beneficial than monoculture AMF for the taller species *G. mollugo* and *L. pratensis*, but not for the three shorter species *P. lanceolata, P. vulgaris* and *V. chamaedrys.* All studied plant species except *V. chamaedrys* showed increased biomass production in the presence of AMF; and across species, plant biomass generally increased with increasing AMF colonization. Interestingly, mixture AMF showed lower colonization than monoculture AMF in the plant species that benefited from them and vice versa in the other three species. These species-specific responses are in line with other studies showing context-dependent AMF effects on plants (Burrows & Pfleger, 2002; Hoeksema et al., 2010). Although AMF generally promote plant growth, the outcome of the interaction may vary from beneficial to antagonistic for plant growth (Johnson, Graham, & Smith, 1997; Kiers & van der Heijden, 2006; Klironomos, 2003). The positive effect of mixture AMF on species with a low colonization could indicate increased AMF efficiency in promoting plant growth. However, lower growth at high colonization could also indicate that a shift from mutualism to antagonism occurred in these species.

Colonization in our positive AMF control treatment of *R. irregulare*, which did not share a common history with the experimental plants, was greater than in mixture or monoculture AMF treatments and led to higher plant biomass especially at the first harvest. A high colonization ability of this taxon is typical and has been shown to be not as beneficial to plant hosts as less abundant colonization by other AMF taxa (Hart and Reader 2002, Wagg et al., 2011a, Engelmoer, Behm, & Kiers 2014). This particularly productive and cosmopolitan AMF taxon may have been more effective in colonizing plant roots and influencing plant responses compared to the inoculation of AMF spore communities native to the field site where a greater diversity of AMF taxa were likely present (Dassen et al., 2017). The coinoculation of several AMF taxa could have resulted in competition among AMF taxa for a single host plant in our study, thus reducing their ability to colonize roots and influence plant responses (Bennett & Bever, 2009; Engelmoer et al., 2014). These results are in line with previous findings reporting that different AMF isolates vary in their effect on plant phenotypes (Klironomos, 2003; Koch, Antunes, Maherali, Hart, & Klironomos, 2017; Streitwolf-Engel, Boller, Wiemken, & Sanders, 1997), which may depend on co-infection with other AMF species (Arguello et al., 2016; Bennett & Bever, 2009).

### 4.2 INFLUENCE OF PLANT DIVERSITY HISTORY ON PLANT PHENOTYPES (H2)

Our second hypothesis, that mixture-type plants grow faster whereas monoculture-type plants are better defended was generally supported by our experiment (see significant main effects of plant history in Table 1). But here again the plant species varied in their responses. Mixture-type plants of three species were more productive and mixture-type plants of three species grew taller than monoculture-type plants. Furthermore, three out of the five species showed higher LDMC and four species higher LMA for mixture-type plants in comparison with monoculture-type plants, indicating that mixture-type plants also invested more resources into leaf biomass production. In contrast, monoculture-type plants had less leaf damage, which confirmed the second part of our hypothesis 2, namely that monoculture-type plants evolved increased defence. We observed particularly severe infection by powdery mildew in mixture-type plants of *P. lanceolata*, suggesting, in agreement with Engelmoer et al. (2014), that monoculture-type plants of *P. lanceolata* may have been subjected to particularly strong selection pressure for pathogen defence in comparison with mixture-type plants. Previously it has been shown that community diversity as selective environment can alter the selection on plant traits favouring greater character displacement in more diverse plant communities (Zuppinger-Dingley et al., 2014), with the assumption that competitive growth traits trade off with selection for pathogen defence in plant monocultures (Herms & Mattson, 1992). The increased resource investment into leaves in mixture-type plants thus may have been at the cost of lowered pathogen defence. Specialized pathogens have indeed been observed to become diluted at higher plant diversity levels in the Jena Experiment (Eisenhauer, Reich, & Scheu, 2012; Rottstock, Joshi, Kummer, & Fischer, 2014), thus conferring lower selection pressure in mixtures than in monocultures. To increase survival, plants in monocultures are expected to allocate more resources to defence (Bezemer & Vandam, 2005) or may enter more beneficial symbioses (Newsham et al., 1995). Consequently, survival in plant monocultures may depend on the ability of plants to allocate resources to these interactions whereas plants in mixtures can allocate these resources to growth.

### 4.3 HOME VS. AWAY EFFECTS OF PLANT AND AMF HISTORIES (H3)

Our third hypothesis was that a common selection history of plants and their associated fungi in the field would lead to a matching between plants and their associated AMF communities, i.e. ‘home’ vs. ‘away’ AMF effects on plant phenotypic responses. We found limited evidence for this hypothesis. The main predictor for increased positive PSF, plant biomass, was not affected by home vs. away combination, which indicates that beneficial interactions did not strengthen over time. Instead, we found a home vs. away effect for the two functional traits LMA and leaf absorbance. For these two leaf properties, we found some evidence that monoculture-type plants were more influenced by monoculture AMF and mixture-type plants were more influenced by mixture AMF. Namely, leaf absorbance and LMA were increased in the small herb *P. vulgaris* if the AMF had a common selection history. Increased leaf absorbance and high LMA are related to higher area-based nitrogen content (Niinemets, 1997), suggesting that co-selected AMF may have improved the nitrogen uptake of these plants. The opposite response was observed for leaf absorbance in *L. pratensis* and for LMA in *V. chamaedrys.* Although for LMA and leaf absorbance the results would be in line with our hypothesis for a specific effect of ‘matching’ plant and AMF community histories (indicating their co-selection), the common selection history between plants and AMF was rather detrimental to the plants’ growth response. This would suggest that a common association history between plants and their AMF communities may have become more antagonistic, rather than mutualistic, over the eleven years of co-selection.

## 5.1 Conclusions

Here we present evidence that the loss of plant diversity differentially alters how AMF communities influence phenotypic responses in plants. We found that AMF communities selected in plant mixtures were more beneficial than AMF selected in plant monocultures for two out of five plant species. Furthermore, mixture-type plants generally grew better, but suffered more leaf damage than monoculture-type plants, providing evidence for the differential selection on competitive growth vs. defence depending on the interspecific diversity of neighbouring plants. However, home vs. away effects between AMF and plants from mixtures or AMF and plants from monocultures were rare. When they did occur they were generally in the direction of increased antagonism by the AMF, leading to a reduced performance of the plant partner. We assert that conducting reciprocal inoculation experiments over longer ecological time-scales could contribute to a better understanding of plant–AMF interactions (Lekberg & Koide, 2014). Our results suggest that biodiversity loss can alter evolutionary trajectories in the interactions between plants and AMF through differential selection pressures on both partners at high and low diversity. Furthermore, our data suggest that a shared history between plants and AMF does not follow a general pattern leading to increased mutualism. Rather, changes in these interactions are context-dependent and may strengthen over time, even switching directionality. Further investigating the sustainability of AMF-plant mutualisms in a world with changing species composition and biodiversity will be important when searching for applications for AMF communities to support ecosystem functioning in the future.

## Acknowledgements

We thank M. Furler, D. Topalovic, D. Trujillo Villegas and T. Zwimpfer for technical assistance and A. Ferrari, S. Karbin, J. Moser Frias and Y. Xu for help with measurements. Thanks to M. van der Heijden for assistance and the AMF isolates. This study was supported by the Swiss National Science Foundation (grants number 147092 and 166457 to B. S.) and the University Research Priority Program Global Change and Biodiversity of the University of Zurich. The Jena Experiment is supported by the German Science Foundation (FOR 1451, SCHM 1628/5-2).

## Authors’ contributions

T.H., C.W., S.J.V.M., D.Z.D and B.S. planned and designed the research, T.H. carried out the experiment and T.H., C.W., M.W.S., S.J.V.M and B.S. analyzed the data. S.J.V.M, T.H., C.W. and B.S. wrote the manuscript with the other authors contributing to revisions.

## Data accessibility

Data will be archived on Pangaea upon acceptance of this article.

## Supporting information

Additional Supporting Information may be found online in the Supporting Information tab for this article:

**Methods S1**. Assessment of soil N and P content at the beginning of the experiment.

**Fig. S1** Aboveground biomass production of monoculture- and mixture-type plants in control soil in dependence of different levels of AMF colonization.

**Table S1.** Experimental design.

**Table S2.** Plant traits measured at the experimental plant age.

